# Visualizing single-cell data with the neighbor embedding spectrum

**DOI:** 10.1101/2024.04.26.590867

**Authors:** Sebastian Damrich, Manuel V. Klockow, Philipp Berens, Fred A. Hamprecht, Dmitry Kobak

**Affiliations:** Hertie Institute for AI in Brain Health, University of Tübingen, Germany; Tübingen AI Center, Tübingen, Germany; IWR, Heidelberg University, Germany; Konrad Zuse School of Excellence in Learning and Intelligent Systems (ELIZA)

## Abstract

The two-dimensional embedding methods *t*-SNE and UMAP are ubiquitously used for visualizing single-cell data. Recent theoretical research in machine learning has shown that, despite their very different formulation and implementation, *t*-SNE and UMAP are closely connected, and a single parameter suffices to interpolate between them. This leads to a whole spectrum of visualization methods that focus on different aspects of the data. Along the spectrum, this focus changes from representing local structures to representing continuous ones. In single-cell context, this leads to a trade-off between highlighting rare cell types or continuous variation, such as developmental trajectories. Visualizing the entire spectrum as an animation can provide a more nuanced understanding of the high-dimensional dataset than individual visualizations with either *t*-SNE or UMAP.

---

Single-cell transcriptomic datasets can contain gene expression measurements of thousands of genes in millions of cells (Svensson et al., 2020). Such high-dimensional datasets are notoriously difficult to analyze and to interpret. Many statisticians have argued that “often the most effective way to describe, explore, and summarize a set of numbers – even a very large set – is to look at pictures of those numbers” (Tufte, 2001, Introduction), as the human visual system excels at pattern recognition (Kaas and Balaram, 2014). Indeed, single-cell biologists regularly use two-dimensional visualizations for data exploration and hypothesis generation. The two most popular algorithms are neighbor embedding methods *t*-SNE (van der Maaten and Hinton, 2008) and UMAP (McInnes et al., 2018).

Such 2D embeddings are necessarily distorted (Wattenberg et al., 2016; Chari and Pachter, 2023; Wang et al., 2023b): they cannot faithfully represent high-dimensional distances, and so cluster sizes, densities, and relative positions cannot be directly interpreted. Moreover, humans sometimes recognize patterns *too* well (Kahneman and Tversky, 1972; Gilovich, 2008) and can over-interpret even random noise: “to the untrained eye randomness appears as regularity or tendency to cluster” (Feller, 1991, p.161). On the other hand, multiple benchmark studies have shown that *t*-SNE and UMAP embeddings do capture some biologically relevant information, such as cluster structure or transitions between related cell types (Espadoto et al., 2021; Sun et al., 2019; Xiang et al., 2021; Xia et al., 2021; Huang et al., 2022; Wang et al., 2023a; Lause et al., 2024), and many biologists find them useful when working with single-cell data.

The exact relationship between UMAP and *t*-SNE has long been unclear and has generated some controversy (Becht et al., 2019; Kobak and Linderman, 2021), making it difficult for practitioners to make an informed choice about when to use which method. Visually, UMAP embeddings often feature more condensed and clear-cut clusters than *t*-SNE (Böhm et al., 2022), which may have been one of the reasons for its fast adoption by the singlecell community.

We have recently described a clear conceptual relation between *t*-SNE and UMAP and hence shown that they can be transformed into each other by varying a single hyperparameter (Böhm et al., 2022; Damrich et al., 2023). Here we argue that this hyper-parameter is highly relevant for single-cell applications, as it controls the trade-off between representing local and continuous structures.

All neighbor embedding methods first find nearest neighbors of each individual cell in the high-dimensional data. Then the cells are arranged in two dimensions driven by attraction between nearest neighbors and by repulsion between all cells. The key difference between *t*-SNE and UMAP is how they balance attraction and repulsion: *t*-SNE uses weaker attraction than UMAP. All other differences only have minimal impact on the embedding (Böhm et al., 2022; Damrich and Hamprecht, 2021; Damrich et al., 2023).

Varying the relative strength of attraction and repulsion leads to a whole *neighbor embedding spectrum*. This allows to interpolate between *t*-SNE and UMAP, but also to achieve embeddings beyond *t*-SNE or UMAP. Higher attraction levels lead to embeddings similar to ForceAtlas2 (Böhm et al., 2022) and eventually to Laplacian Eigenmaps (Linderman and Steinerberger, 2019).

Different embeddings on the spectrum represent different types of structure better. High attraction strongly pulls similar cells together, resulting in condensed clusters and more pronounced transitions between adjacent clusters. This often makes continuous structures such as developmental trajectories more visible. Conversely, with high repulsion, finer cell types are able to drift further apart in the embedding, improving the local arrangement of cells.

Indeed, empirical benchmarks found that *t*-SNE typically performs best in local metrics quantifying neighborhood preservation (Xiang et al., 2021; Huang et al., 2022; Wang et al., 2023a; Lause et al., 2024), while UMAP and ForceAtlas2 can perform better in global metrics such as correlation between pairwise distances (Huang et al., 2022; Sun et al., 2019), in particular for developmental data with pronounced continuous structure.

We illustrate this using data from human brain organoid development (Kanton et al., 2019). High-repulsion embeddings, such as *t*-SNE (Figure 1, left), show detailed local structure (inset), whereas high-attraction embeddings (Figure 1, right) make the developmental trajectory (dashed line) more prominent. This visual impression is confirmed by quantitative metrics showing the trade-off between local and global structure preservation (Figure 2).

**Figure 1:**
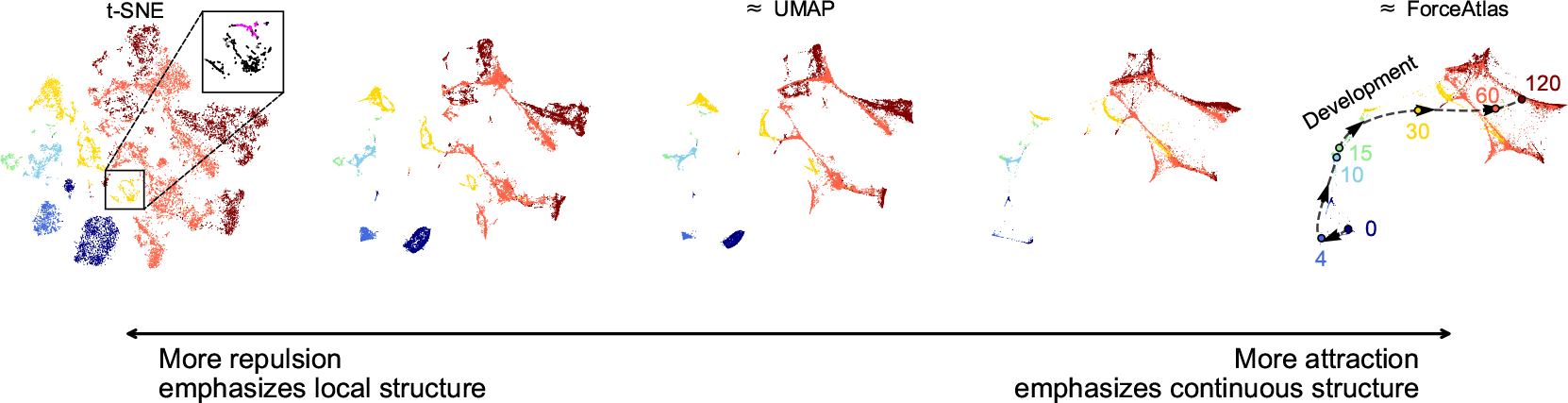
Neighbor embedding spectrum. Brain organoid development data (Kanton et al., 2019) colored by the sample age (from 0 to 120 days). Sample size *n* = 20 272. Stronger repulsion shows finer local details. For instance, the inner structure of the island in the enlarged inset corresponds to two different cell types (colored magenta and black) identified by Kanton et al. (2019). Conversely, stronger attraction brings out the developmental trajectory (dashed line). *t*-SNE, UMAP, and ForceAtlas2 lie on this spectrum. Embeddings were computed via `openTSNE` with exaggeration values 1, 2.5, 5, 15, 30.

**Figure 2:**
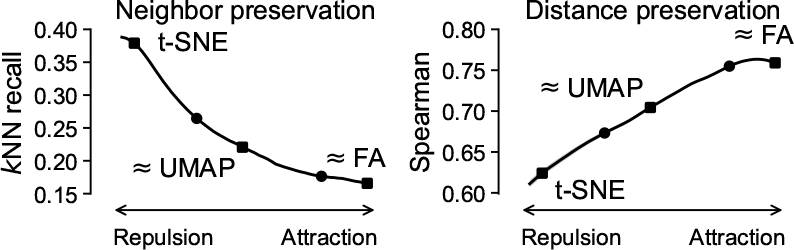
Local / global trade-off. along the neighbor embedding spectrum of the Kanton et al. (2019) data. Neighbor preservation, a local metric, is measured via *k*NN recall (*k* = 15). Distance preservation, a global metric, is measured via the Spearman correlation between pairwise distances. Points correspond to panels of Figure 1. We plot mean *±* one standard deviation across three random seeds.

In practical single-cell applications, higher repulsion may be more appropriate for visualizing atlases of fully differentiated cells, whereas higher attraction can be suitable for developmental datasets. That said, we suggest to look at multiple embeddings along the spectrum, paying attention to which structures persist and which vanish. This is similar to interactive exploration strategies common in wet-lab biology like changing staining intensity or microscope lens, and can give more insight into the structure of the high-dimensional data than any given isolated embedding. In particular, it can be convenient to visualize the spectrum as an animation with smooth transitions between the attraction parameter values (Suppl. File).

Practically, the attraction strength can be varied both in a *t*-SNE-based loss function and in sampling-based losses similar to UMAP’s loss. This functionality is available in various existing implementations such as openTSNE (Poličar et al., 2019) or CNE (Damrich et al., 2023). Figures 1,2 used openTSNE. Appendix A contains the analogous figures based on CNE. We provide a Python package ne-spectrum for computing the whole neighbor embedding spectrum and for making animations. Our package relies on openTSNE and CNE libraries to compute individual embeddings, starting with high attraction and gradually decreasing the attraction strength. The whole process only takes 2–5 times longer than computing a single embedding.

In conclusion, *t*-SNE, UMAP, ForceAtlas2, and Laplacian Eigenmaps correspond to different settings on the attraction-repulsion spectrum of neighbor embeddings. Different settings on the spectrum emphasize different kinds of biological structure and no single setting is optimal for all purposes. Therefore, we recommend to use the whole spectrum for data exploration, for instance in form of an animation, instead of relying too much on any individual embedding.

Practitioners should be aware that 2D embeddings can strongly distort high-dimensional data and carry the risk of being over-interpreted (‘seeing is believing’). Looking at the entire spectrum mitigates the risk of jumping to premature conclusions based on any given 2D picture of high-dimensional data. More generally, we argue that the proper use of 2D embeddings is to suggest and guide further computational analysis. Once findings are confirmed, the spectrum can help finding the individual visualization most appropriate for scientific communication, where it should be treated as a suggestive illustration.

## Supporting information

Animated NE spectrum via CNE

Animated NE spectrum via openTSNE

## Acknowledgements

This work was funded by Deutsche Forschungsgemeinschaft (DFG, German Research Foundation) via SFB 1129 (Projektnummer 240245660) as well as via Germany’s Excellence Strategy (Excellence cluster 2064 “Machine Learning — New Perspectives for Science”, EXC 390727645; Excellence cluster 2181 “STRUCTURES”, EXC 390900948), the German Ministry of Science and Education (BMBF) via the Tübingen AI Center (01IS18039A) as well as via the DAAD programme Konrad Zuse School of Excellence in Learning and Intelligent Systems (ELIZA), the Gemeinnützige Hertie-Stiftung, and the National Institutes of Health (UM1MH130981). The content is solely the responsibility of the authors and does not necessarily represent the official views of the National Institutes of Health.

## Data and code availability

We used the data from Kanton et al. (2019) taken from ArrayExpress with the accession code E-MTAB-7552. Our code for reproducing the figures, including scripts for downloading and preprocessing the data, is available at https://github.com/berenslab/ne_spectrum_scRNAseq. The package for computing the spectra is available at https://github.com/sciai-lab/ne-spectrum and can be installed with pip install ne-spectrum. The relevant commits in both repositories are tagged bioRxiv-v1.

## A Supplementary Figures

**Figure S1:**
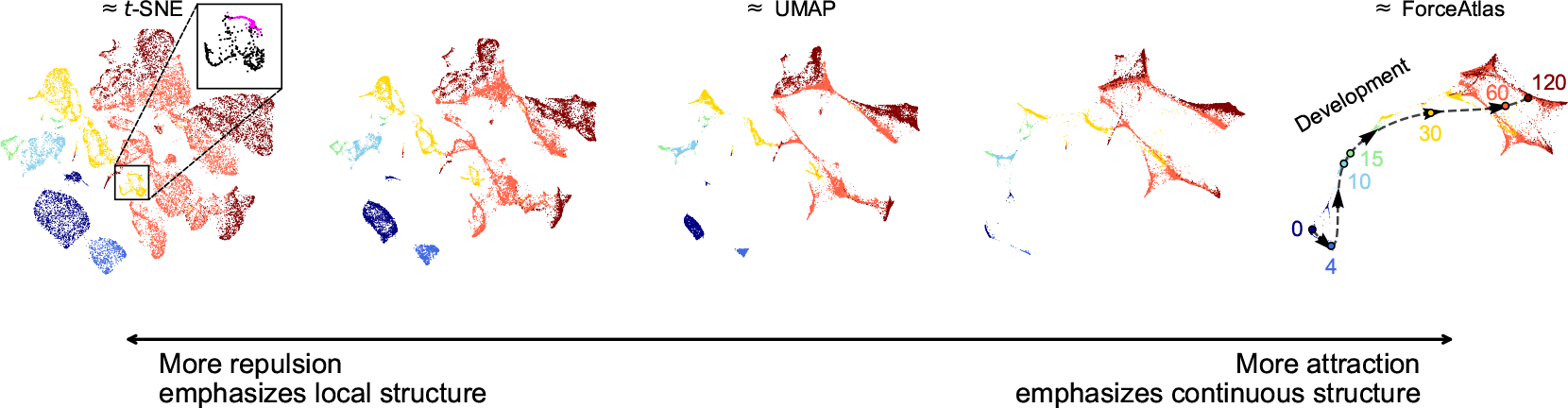
Neighbor embedding spectrum computed with a contrastive loss. Brain organoid development data (Kanton et al., 2019) colored by the sample age (from 0 to 120 days). Sample size *n* = 20 272. Stronger repulsion shows finer local details. For instance, the inner structure of the island in the enlarged inset corresponds to two different cell types (colored magenta and black) identified by Kanton et al. (2019). Conversely, stronger attraction brings out the developmental trajectory (dashed line). *t*-SNE, UMAP, and ForceAtlas2 lie on this spectrum. Embeddings were computed via `contrastive-ne` with s values 0.0, 0.5, 1.0, 1.5, 2.0 and the negative sampling loss using 50 negative samples. We used 50 negative samples instead of 5, which is default in CNE and UMAP, for better neighbor preservation in the high-repulsion setting.

**Figure S2:**
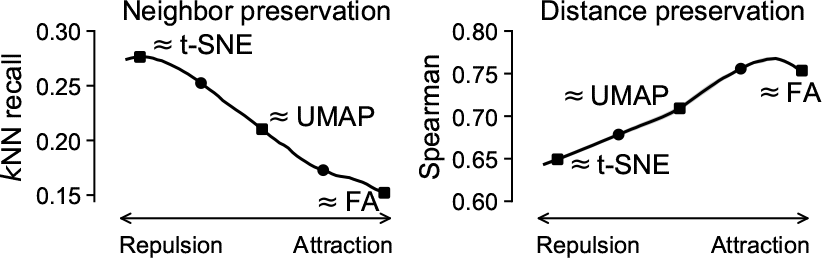
Local / global trade-off. along the neighbor embedding spectrum of the Kanton et al. (2019) data. Neighbor preservation, a local metric, is measured via *k*NN recall (*k* = 15). Distance preservation, a global metric, is measured via the Spearman correlation between pairwise distances. Points correspond to panels of Figure S1. We plot mean *±* one standard deviation across three random seeds. We used 50 negative samples to achieve better neighbor preservation than with default 5 negative samples. While the overall level of neighbor preservation was still lower than in Fig. 2, it can be further increased by using even more negative samples (Damrich et al., 2023, Figs. S18, S19). The Spearman correlation is similar to that of the `openTSNE` backend in Fig. 2.

