## Supplementary figures and images for "Visualizing single-cell data with the neighbor embedding spectrum"

### Animated NE spectrum via CNE

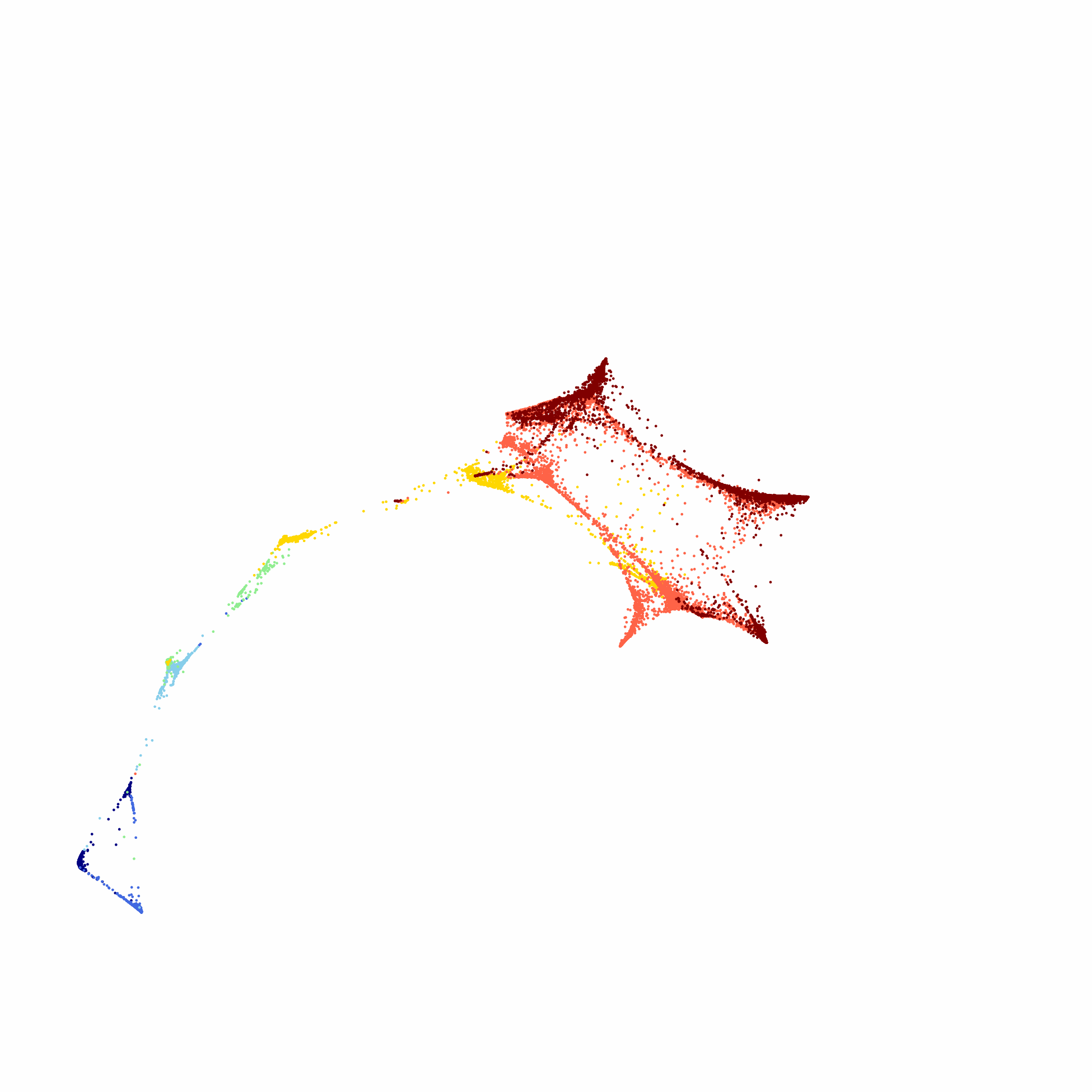

### Animated NE spectrum via openTSNE

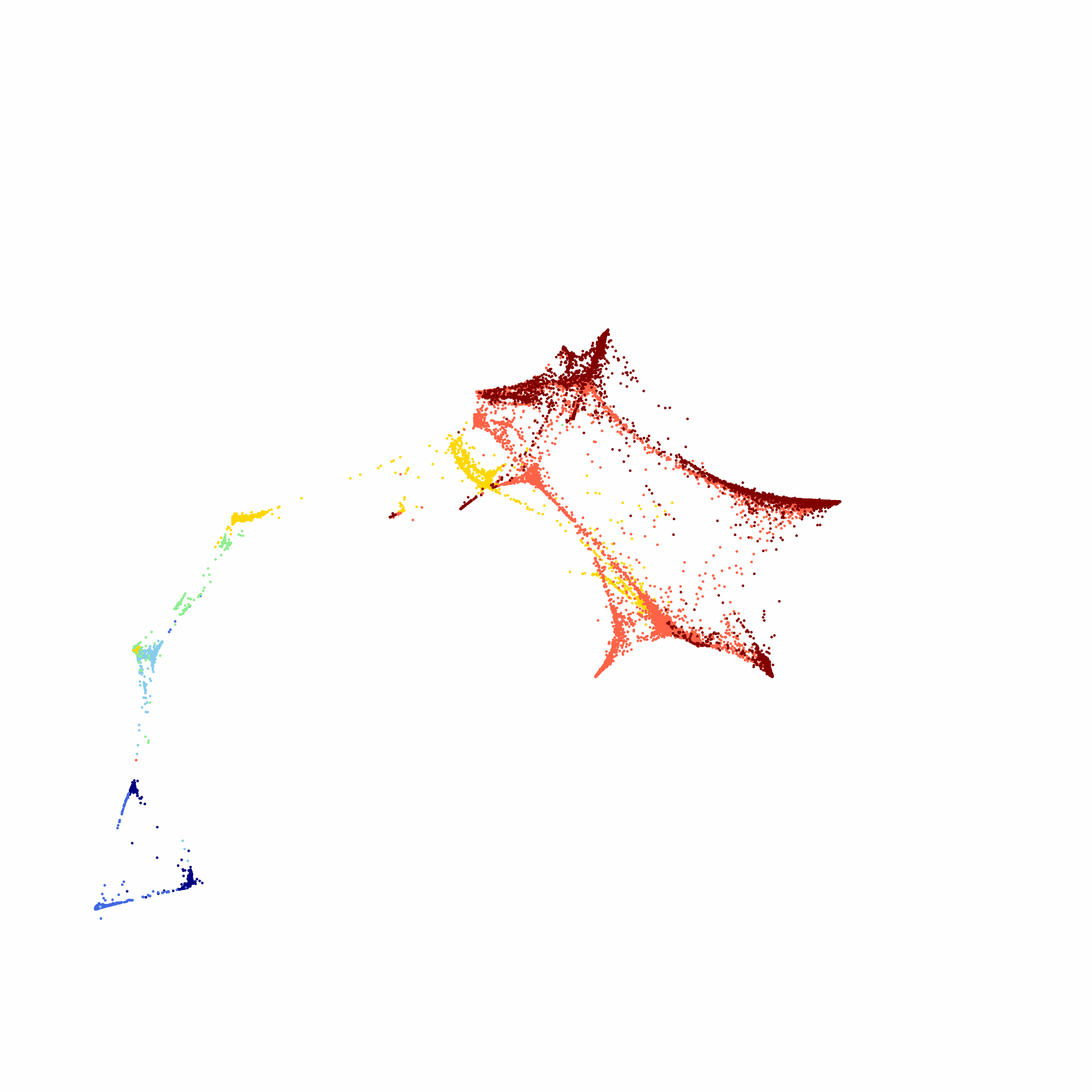
